# The sea urchin intestinal microbiome responds dynamically to food intake and contains nitrogen-fixing symbionts

**DOI:** 10.1101/2024.02.25.581913

**Authors:** Mia M. Bengtsson, Marita Helgesen, Haitao Wang, Stein Fredriksen, Kjell Magnus Norderhaug

## Abstract

Kelp deforestation by sea urchin grazing is a widespread phenomenon globally, with vast consequences for coastal ecosystems. The ability of sea urchins to survive on a kelp diet of poor nutritional quality is not well understood and bacterial communities in the sea urchin intestine may play an important role in digestion. A no-choice feeding experiment was conducted with the sea urchin *Strongylocentrotus droebachiensis*, offering three different seaweeds as diet, including the kelp *Saccharina latissima*. Starved sea urchins served as experimental control. Amplicons of the 16S rRNA gene were analyzed from fecal pellets. One dominant symbiont (*Psychromonas marina*) accounted for 44 % of all sequence reads and was especially abundant in the sea urchins fed seaweed diets. The starved and field captured sea urchins consistently displayed higher diversity than the seaweed-fed sea urchins and displayed a higher predicted abundance of genes involved in nitrogen fixation. Cloning and sequencing of the NifH gene revealed diverse nitrogen fixers. We demonstrate that the sea urchin intestinal microbiome is dynamic, responds to diet and has the capacity for nitrogen fixation. The microbiome thereby reflects the dietary flexibility of these sea urchins, and could be a key component in understanding catastrophic kelp forest grazing events.

## INTRODUCTION

Sea urchins are key stone species in coastal ecosystems and a prominent grazer transforming highly productive kelp forests into unproductive marine deserts (Filbee-Dexter and Scheibling 2014). In the past century, kelp forests in many parts of the world have been turned into so-called “barren grounds” due to excessive grazing by sea urchins (Bernstein, Williams and Mann 1981; Scheibling, (null) and Balch 1999; Norderhaug and Christie 2009). For example, grazing by the sea urchin *Strongylocentrotus droebachiensis* led to the loss of 2 000 km^2^ of *Laminaria hyperborea* and *Saccharina latissima* kelp forest in northern Norway from the 1970s until today (Norderhaug *et al*. 2021). The barren ecosystem can persist for decades, due to constant grazing by urchins, which prevents the regrowth of kelp. When the kelp forest ecosystem is transformed into barren ground, a significant loss in biodiversity and productivity will follow, and numerous important ecosystem services will be lost (Eger *et al*. 2023). One ecosystem service that has received increasing attention during the last decade is the role of kelp forests in climate mitigation by its high efficiency in capturing CO2 and what is referred to as “blue carbon” (Krause-Jensen *et al*. 2018). Large quantities of fragmented and dead kelp detritus are transported to deeper water, buried in sediments and thereby removed from the carbon cycle. Urchins have a significant role in this carbon sink function. They can consume a large part of the drift kelp on shallow water (Filbee-Dexter *et al*. 2019), and by transforming drift kelp into urchin pellets, increase the dispersal potential and extend carbon export into deep water (Wernberg and Filbee-Dexter 2018).

Sea urchins are omnivores as they consume a broad diversity of organisms from several trophic levels, yet, the primary food source is seaweed like kelp (Himmelman and Steele 1971). The ability to live on fresh kelp tissue is rare among animals, and few organisms other than sea urchins can live directly from fresh kelp (Mann 1977). Kelp is difficult to digest because of complex carbohydrates such as alginate that are not readily decomposed by animal enzymes (Lasker and Giese 1954), and the lack of protein. This results in a C:N ratio (Carbon and Nitrogen ratio) exceeding what is found in most marine organisms, which means that kelp has poor nutritional quality (Sterner and Hessen 1994; Norderhaug, Fredriksen and Nygaard 2003).

In order to survive on such an unpalatable diet, sea urchins require adaptations to digest complex carbohydrates and an ability to compensate for the lack of proteins. Other organisms that live on carbon-rich materials can give some indications to how sea urchins can survive on a kelp-dominated diet. For example, shipworms have established symbiotic relationships with bacteria in their guts that makes it possible to live on wooden materials (Lechene *et al*. 2007; Betcher *et al*. 2012). Some of these bacteria decompose complex carbohydrates and others fix nitrogen into biologically available ammonia (NH3), thus compensating for the lack of protein in the food (Lechene *et al*. 2007). Bacteria with similar functions are likely beneficial for sea urchins as well. In fact, nitrogen fixation has been detected in association with some sea urchins, including *S. droebachiensis* (Guerinot and Patriquin 1981a) and a nitrogen-fixing *Vibrio* sp. strain was isolated in culture from the same species (Guerinot and Patriquin 1981b). Despite these early findings, the possible role of the sea urchin microbiome in catastrophic kelp forest grazing events has received relatively little attention. One recent study investigated the intestinal microbiome of *Strongylocentrotus purpuratus* and found that it is highly variable between gut tissue and gut digesta (Hakim *et al*. 2019), indicating a spatial separation in microbial niches in the gut. The intestinal microbiome of *S. purpuratus* was also found to be distinct from another co-occurring sea urchin species, and varied depending on whether it was sampled from kelp beds or from barren grounds (Miller *et al*. 2021). Similarly, the gut microbiome of another sea urchin, *Mesocentrotus nudus*, differed depending on barren ground severity (Park *et al*. 2023). *M. nudus, S. purpuratus* and *S. droebachiensis* all cause catastrophic grazing of kelp forests, and the latter is a quantitatively relevant grazer of kelp in the entire northern hemisphere (Filbee-Dexter and Scheibling 2014).

In this study we set out to fill critical knowledge gaps in the understanding of sea urchin grazing by investigating the intestinal microbiome of *S. droebachiensis* (green sea urchin) during grazing and digestion of seaweeds. We experimentally exposed sea urchins to different seaweed diets and hypothesized that 1) different seaweed as a food source lead to different intestinal microbiome composition during food degradation and that 2) bacterial N2 fixing symbionts are present in sea urchin intestines. We used 16S rRNA gene amplicon sequencing to study differences in bacterial community composition and diversity between different food sources and to predict microbial functions. Cloning and sequencing of the NifH gene was used to detect and identify potential nitrogen-fixing members of the microbiome.

## MATERIALS & METHODS

### Sea urchin collection and experimental design

Sea urchins (*Strongylocentrotus droebachiensis* O.F. Müller) approximately 40-60 mm in diameter were collected next to Hallangstangen (59°40’58.8”N, 10°36’49.0”E, Figure 1), 4 km north of the main harbor in Drøbak (Norway) on January 4th, 2017. As sea urchins are mostly found on hard substrata where they can be firmly attached to the surface, a triangular dredge was used to collect the sea urchins from 6-15 m depth.

**Fig. 1:**
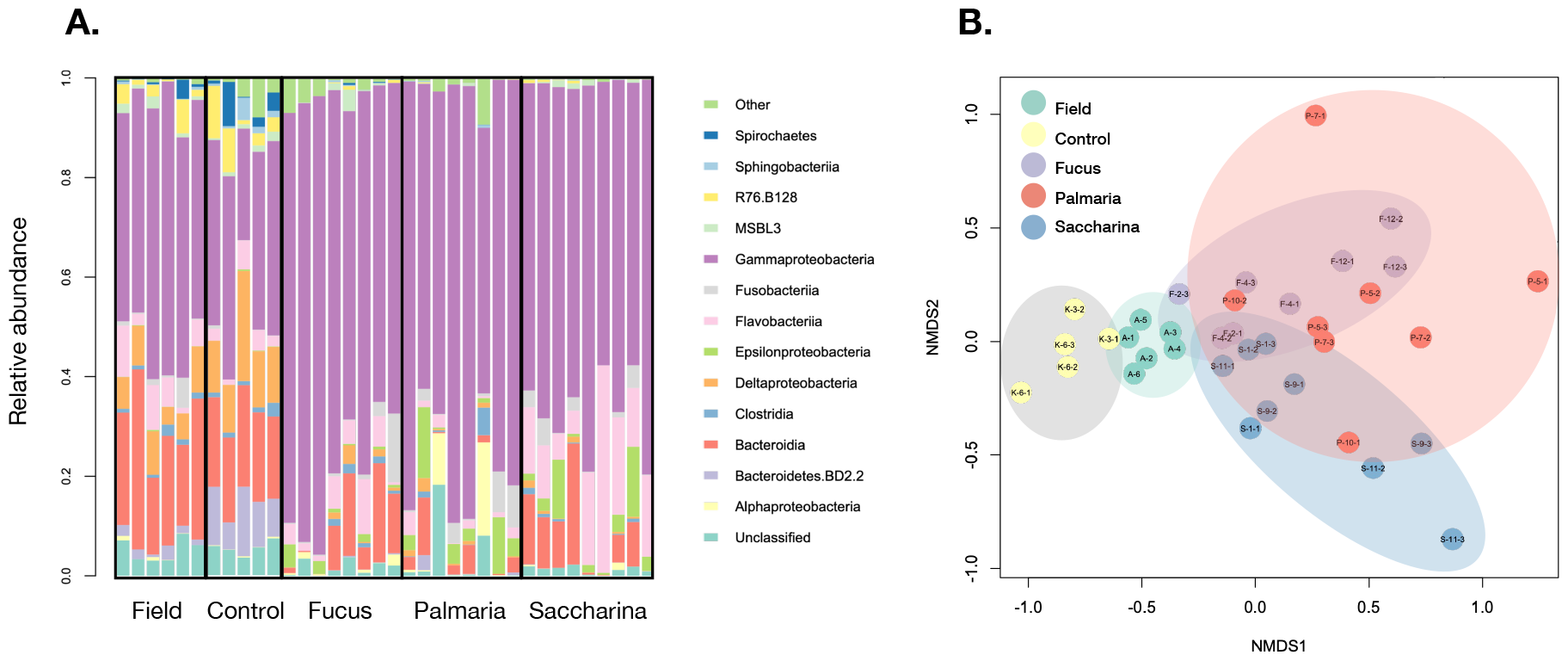
Sea urchin intestinal microbiome composition responded to food availability. (A) Gammaproteobacteria dominated most samples, especially in the treatments where urchins were fed seaweed (seaweed-fed: *Fucus, Palmaria, Saccharina*). (B) The nMDS plot shows a separation between starved (Control) and field-collected (natural diet control) vs. seaweed-fed treatments. The shaded ellipses have been added to aid in interpretation and do not represent confidence intervals.

For the experiment, sea urchins were randomly distributed among 12 tanks (L x W x H: 35 x 20 x 25 cm = 17.5 L) with five individuals in each tank (5 urchins x 12 tanks = 60 urchins). The set-up was designed as a flow-through system, where each tank had its own filter transporting new seawater from a large container, that was continuously re-filled from an inlet next to the Biological station in Drøbak. This design was selected to prevent mixing of water among tanks, and to provide natural water conditions for the sea urchins. The rate of water flow was between 33 to 67 L/h. The urchins were kept in the tanks and starved for ten days prior to implementing the experimental treatments, to minimize the influence of previous feeding and to let them acclimatize to the new conditions. Water temperature (ranging between 3.5-5.4°C) and salinity (26.2 - 30.3 ppt) were monitored on a regular basis to assure good conditions for the sea urchins. The lights were turned off during the experiment (except when handling), to reduce the impact of undesirable growth by algae in the tanks.

### Experimental treatments

A no-choice feeding experiment was conducted to determine the influence of diet on the intestinal microbiome. The diets consisted of three seaweed species: *Saccharina latissima* (Linnaeus) C.E. Lane, C. Mayes, Druehl & G.W. Saunders, *Fucus serratus* Linnaeus and *Palmaria palmata* (Linnaeus) F. Weber & D. Mohr. These algae were selected as they were present in the area where the sea urchins were collected. *S. latissima* (treatment “Saccharina”) is the dominating kelp on sheltered coasts, and the other two algae (treatments “*Fucus*” and “*Palmaria*”) were selected as they generally have different nutrient and chemical profiles compared to kelp, and can function as an alternative food source when kelp is absent. Control tanks (treatment “Control”) with sea urchins which were not fed (starved) were also set up to evaluate if the experimental conditions impacted the results. In addition, samples were taken directly from the collected sea urchins at the field site before initiating the experiment, to represent a natural microbiome (treatment referred to as “Field”). The seaweed used as feed was sampled prior to the experiment to estimate the carbon and nitrogen content using an elemental analyzer (Thermo Finnigan).

### Intestinal microbiome sample collection

After being exposed to the experimental treatments for 10 days, sea urchin intestinal microbiomes were assessed by collection of fecal pellets directly from dissected sea urchins. Although invasive, this method was preferred over collecting fecal pellets in the experimental tanks, to avoid contamination and degradation of fecal material by the surrounding water, and to assign the sampled material to distinct individuals. Fecal pellet samples were taken from 3 urchins from each experimental tank (3 urchins * 12 tanks = 36 samples), in addition, samples were taken from urchins before the experiment (6 individuals, “field” treatment). The sea urchins were dissected and fecal pellets from the large intestine were placed into separate cryovials and then frozen directly with liquid nitrogen to prevent DNA from degrading. Between each dissection, the equipment was sterilized with decanox, sterilized water and ethanol (70 %). The samples were stored at - 80°C until DNA isolation. Fecal pellets could not be found in all sea urchins, especially the starved (control) urchins, thus the sample size in all groups was not identical.

### DNA extraction and analysis

DNA isolation was performed with the commercial DNeasyPowerSoil® (Qiagen, formerly MoBio Laboratories). The fecal pellet samples were thawed and kept on ice and samples were handled as fast as possible, since pilot experiments had shown that DNA degradation was a critical issue. The DNA isolation was carried out according to the protocol provided by the manufacturer, with some minor changes: Solution C1 was added before the sample, the amount of fecal pellet samples added was between 0.02 to 0.08 g, and half the amount of solution C6 was used to elute the isolated DNA in the final step. The amount of DNA in isolated from each sample was measured with a Qubit 3.0 fluorometer.

PCR and Illumina MiSeq V3 sequencing were carried out by LGC Genomics in Germany. The primers used were: forward: 515F-mod (5’-GTGYCAGCMGCCGCGGTAA-3’) and reverse: 806R-mod (5’-GGACTACNVGGGTWTCTAAT-3’) (Walters *et al*. 2016), targeting the V4 region of the prokaryotic 16S rRNA gene, and linked to custom barcode and adaptor constructs. The sequence data has been uploaded to the European Nucleotide Archive under the accession number PRJEB54963.

### Sequence processing

The DADA2 R package (Callahan *et al*. 2016) was used to quality-filter and assign amplicon sequence variants (ASVs) from the 16S rRNA amplicon sequencing data. Taxonomic classification of the ASVs generated by DADA2 was carried out using the SILVA Incremental Aligner (SINA), version 1.2.11 (Pruesse *et al*. 2007) on the SILVA homepage using the “search and classify” service, and the gene as “SSU” for small subunit rRNA, otherwise default settings were applied.

### Statistical analysis

All statistical analyses were carried out using R (version 3.4.1) (R Development Core Team 2023). ASV richness (S) was calculated using sequence data rarefied to the lowest sequencing depth for a sample (17 650) using functions *rarefy* and *diversity* from the vegan package (Oksanen *et al*. 2016). To test for differences in richness and evenness between treatments, analysis of variance (ANOVA, *aov* function) followed by a Tukey Post-Hoc test (TukeyHSD function) was carried out. Richness data was log transformed to achieve normality.

To analyze the bacterial community composition, multivariate statistics were implemented. ASV data was Hellinger-transformed with the *decostand* function from the vegan package. The transformed values were used as basis for non-metric multidimensional scaling (*metaMDS* function) plot and permutation analyses of variance (PERMANOVA, *adonis2* function). As there is a risk of sea urchins within the same tank having similar community composition, and that there might be an interaction between treatment and tank variables, an interaction term (treatment*tank) was added to the formula for the PERMANOVA.

### Analysis of N2-fixation potential in sea urchin intestines

To assess whether N2-fixing bacteria play a significant role in sea urchin intestines under the current experimental conditions, two approaches were taken. First, the 16S rRNA sequence data was used to predict the abundance of genes involved in N2-fixation using PICRUSt (Langille *et al*. 2013). To generate a PICRUSt compatible OTU table, a “closed-reference” OTU picking method was implemented using QIIME (Caporaso *et al*. 2010). The OTU table was normalized by dividing the known/predicted 16S rRNA gene copy numbers, and then functions based on KEGG Orthologs (KOs) for metagenomes were predicted using the vector of gene counts for each OTU. KOs corresponding to genes involved in N2 fixation were identified using text search, resulting in 15 KOs (K13819, K04488, K02596, K02588, K02586, K02587, K02597, K02595, K02593, K02592, K02591, K02585, K02584, K00531, K00536). Second, we amplified a fragment of the NifH gene from a subset of the samples using the IGK3/DVV PCR primers (Ando *et al*. 2005) and subsequently cloned these fragments into competent *E*.*coli* cells using TOPO TA cloning. Clones were sequenced using Sanger sequencing technology and the sequences were submitted to GenBank under accession numbers OP380433-OP380446. The obtained DNA sequences were translated to protein sequences which were used as query sequences for searching against the NCBI protein database. Reference sequences were chosen from highly similar sequences to each deduced protein sequence. A maximum likelihood phylogenetic tree was built with the deduced and reference sequences using MEGA7 (Kumar, Stecher and Tamura 2016).

## RESULTS

The Illumina amplicon sequencing resulted in a total of 1 275 250 amplicon sequences across all treatments, and after quality filtering, the number was reduced to 1 097 148 sequences. The total number of filtered sequence-reads present in each sample were between 17 650 to 43 870, with a mean of 28 870 reads. A total of 614 prokaryotic amplicon sequence variants (ASVs) were identified. The majority of the sequences belong to the domain Bacteria, with only one ASV classified as Archaea. ASV no. 1 accounted for 44 % of all sequence reads, demonstrating that ASV no.1 is a dominant symbiont. A BLAST search revealed that ASV no. 1 had a 100 % sequence identity with the cultured species *Psychromonas marina (Gammaproteobacteria)*.

Gammaproteobacteria (to 84% consisting of ASV no 1 *Psychromonas marina*) dominated the sea urchin intestinal microbiome in nearly all sampled individuals, yet this dominance was most pronounced in the sea urchins receiving seaweed, regardless of the seaweed species they were fed (Fig.1a). Starved (control treatment) sea urchins and those sampled directly after collecting in the field (field treatment) displayed relatively distinct community composition, well separated from the seaweed-fed treatments (*Fucus, Palmaria, Saccharina*), which were more overlapping (Fig. 1b). Treatment explained 43% of the variation in the community composition, overwhelming the effect of experimental tank (5.8% of variation explained) according to PERMANOVA (Table 1).

**Table 1.** Results of the multivariate permutational analysis (PERMANOVA) of differences in (hellinger transformed) bacterial communities between treatments (Interaction between the variables treatment and tank are inspected. Treatments: Control, Field, *Fucus, Palmaria, Saccharina*. Formula used: table of amplicon sequence variants (ASV) ∼ treatment*tank. Significance level is indicated by the significant codes: 0 ‘***’ 0.001 ‘**’ 0.01 ‘*’ 0.05 ‘.’ 0.1 ‘’ 1.

|  | Df | Sums of<br>Sqs | Mean<br>Sqs | F.<br>Model | R<sup>2</sup> | Pr(>F) | Sign. c. |
| --- | --- | --- | --- | --- | --- | --- | --- |
| treatment | 4 | 2.7252 | 0.681<br>31 | 6.9766 | 0.43443 | 0.001 | *** |
| tank | 1 | 0.3647 | 0.364<br>72 | 3.7347 | 0.05814 | 0.003 | ** |
| treatment:<br>tank | 3 | 0.5464 | 0.182<br>14 | 1.8651 | 0.08711 | 0.015 | * |
| Residuals | 27 | 2.6367 | 0.097<br>66 |  | 0.42032 |  |  |
| Total | 35 | 6.2731 |  |  | 1.00000 |  |  |

Diversity (ASV richness and evenness) was generally lower in the seaweed-fed treatments than in the control and field treatments. The starved sea urchins in the control treatment displayed the highest diversity (Fig 2a-b). This pattern was echoed in the predicted relative abundance of genes involved in N2 fixation, which was also highest in the starved sea urchins (Fig 2c). Richness was significantly lower in the seaweed feed (*Saccharina, Fucus, Palmaria*) treatments compared to the Control treatment (*p*<0.05) while the Field treatment did not differ significantly (*p*>0.05) from the other treatments with regards to richness. Evenness was significantly lower in the seaweed feed treatments compared to both the Control and the Field treatment (*p*<0.05), while these two treatments did not differ from each other. Neither richness nor evenness was significantly different between any of the seaweed feed treatments.

**Fig. 2:**
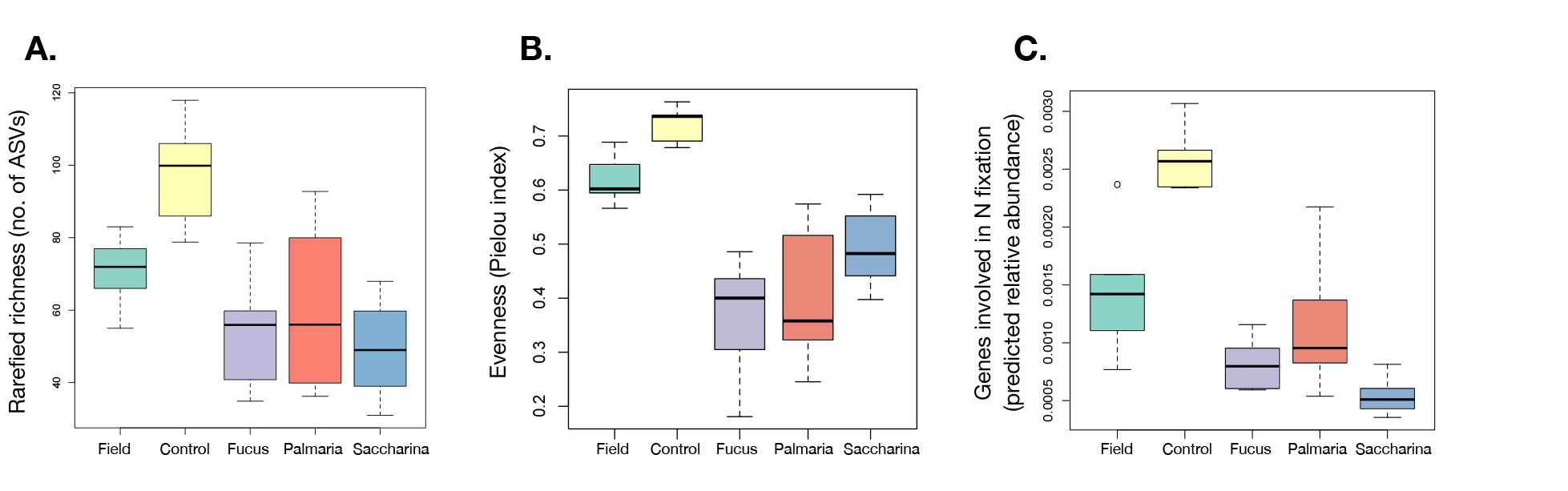
Microbiome diversity and predicted abundance of N_2_ fixation genes were higher in starved (control) and field-collected sea urchins. The starved sea urchins of the control treatment displayed the highest (A) rarefied ASV richness (B) evenness (C) as well as the highest predicted abundance of genes involved in N_2_ fixation (predicted using Picrust on 16S rRNA gene sequencing data).

Amplification and cloning of the *nifH* gene revealed that bacteria capable of N2 fixation were present in the sea urchin intestinal microbiome and belonged to diverse phylogenetic groups (Fig. 3). Most of the successfully amplified and cloned *nifH* variants came from samples from the *Saccharina* treatment, and were related to *Vibrio* spp. (Gammaproteobacteria) from previous studies. Two clones from the Field treatment also fell within this clade. Two clones from the starved control treatment were related to Verrucomicrobia, whereas one clone from the *Saccharina* treatment clustered with *nifH* genes from Bacteroidetes bacteria.

**Fig. 3:**
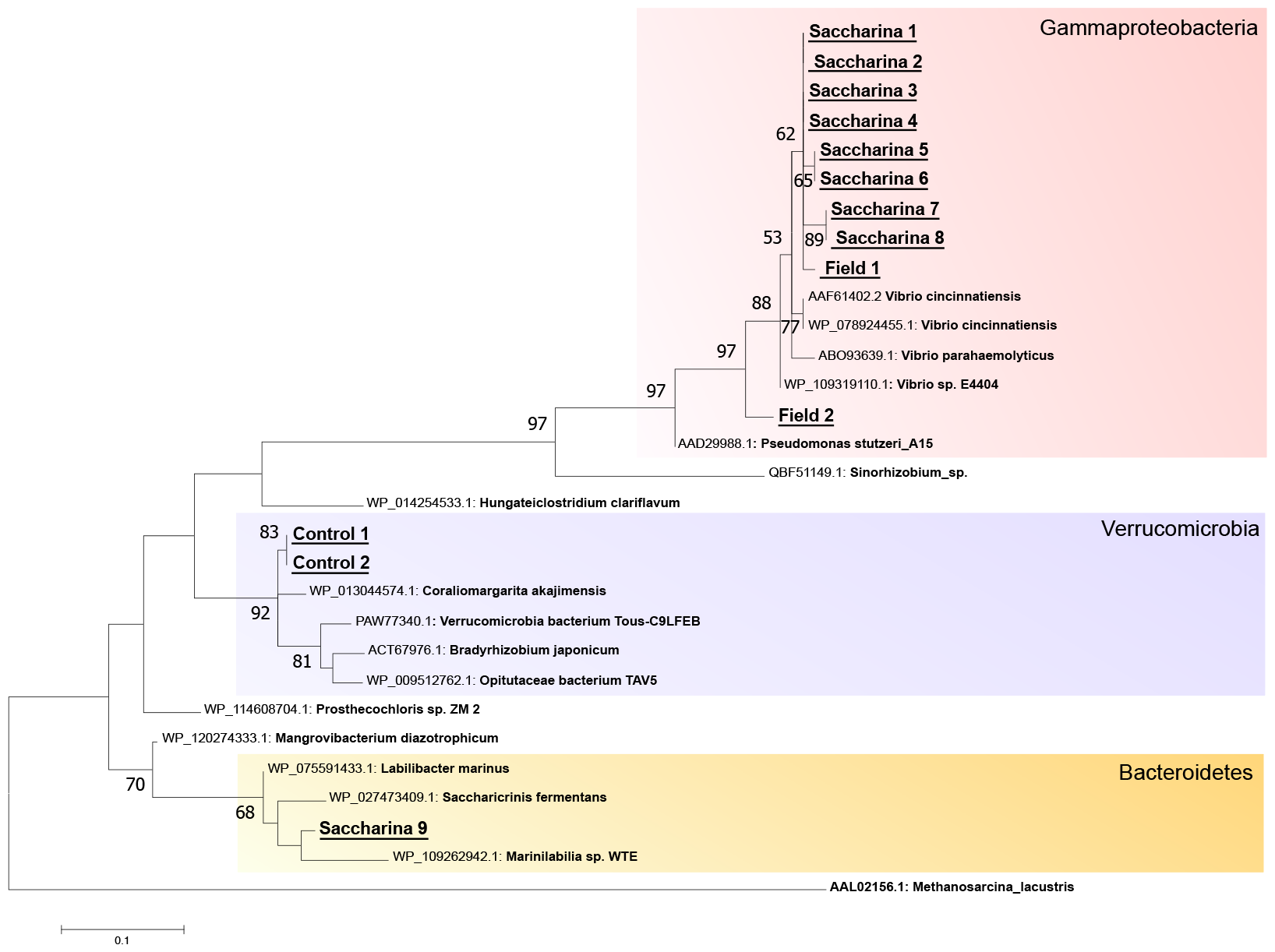
Bacteria capable of N_2_ fixation are present in the sea urchin intestinal microbiome and belong to diverse phylogenetic groups. The Maximum likelihood gene tree was generated using cloned and sequenced fragments of the NifH gene from sea urchins subjected to the control, *Saccharina* (kelp)-fed treatments as well as individuals collected directly from the field.

## DISCUSSION

### Starved and field-captured sea urchins display diverse intestinal microbiomes

This study is the first to assess the intestinal microbiome of the green sea urchin *Strongylocentrotus droebachiensis*, the most important destructive grazer of kelp forests along northern Atlantic coasts (Filbee-Dexter and Scheibling 2014). We found a microbiome dominated by a symbiont closely related to *Psychromonas marina*, yet also displaying a diverse collection of taxa, especially in sea urchins collected directly from the field and those kept starved in aquaria for several weeks. While we hypothesized that different seaweed as a food source would lead to different intestinal microbiome composition during food degradation (hypothesis 1), we rather found that microbiome composition depended primarily on food availability, regardless of food source. The green sea urchin microbiome shared several features with the previously investigated purple sea urchin intestinal microbiome, such as the presence of *Psychromonadaceae* and variability depending on resource availability (Miller *et al*. 2021; Hakim *et al*. 2019). The higher microbial diversity found in starved specimens could indicate the potential for functional plasticity of the microbiome, as a (functionally) diverse microbiome prepares the starved sea urchin for digesting a variety of food types it may encounter. Field-captured sea urchins also featured relatively diverse microbiomes, which may reflect the lower food availability typical for the winter conditions they were sampled from, or a more varied diet compared to our laboratory conditions. In sea urchin barren grounds in South Korea, the severity of the barren ground, which is related to starvation, did not influence gut bacterial diversity although it did influence microbiome composition (Park *et al*. 2023).

### Algal food intake promotes a dominant symbiont related to *Psychromonas marina*

In a feeding experiment, we presented the sea urchins with three different types of seaweed in a no-choice fashion. In all cases, the availability of seaweed led to a decrease in microbial diversity due to an increase in abundance of the dominant symbiont (*Psychromonas ASV1*). *Psychromonas* sp. has also been detected in *S. purpuratus* (Hakim *et al*. 2019; Miller *et al*. 2021) and *S. intermedius* (Zhang *et al*. 2014) and was abundant especially in gut digesta and fecal pellets, analogous to our sample material. Thus, this symbiont appears to be a common feature of the intestinal microbiome of sea urchins belonging to the *Strongylocentrotus* genus, which could point towards important symbiotic functions. Interestingly, several *Psychromonas* OTUs were also detected in association with eggs and larvae of *S. droebachiensis*, and were suggested to play an important role in their development (Carrier and Reitzel 2019). *Psychromonas marina* was originally isolated from cold water off the coast of Japan and is a facultative anaerobe capable of hydrolysing alginic acid, a component of brown algal cell walls (Kawasaki 2002). Different strains of *Psychromonas* sp. have also been described as associated with deep sea amphipods, where their reduced genomes indicate a close symbiotic relationship with their hosts (Zhang *et al*. 2018). The functional capacity of this symbiont in sea urchins remains enigmatic and could not be addressed further with currently available data, yet it is plausible that it is a nutritional symbiont that may assist the sea urchin in degradation and digestion of seaweed biomass. Future investigations should for example uncover genomic or transcriptomic information from the symbiont to elucidate its activity inside the sea urchin intestine or isolate the symbiont in pure culture allowing direct physiological investigation.

### N2 fixing microorganisms were detected in both starved and fed sea urchins

We also hypothesized that nitrogen fixation is an important bacterial symbiont trait during seaweed digestion (hypothesis 2) based on the high C:N ratio of the diet (Table 2) and the previous detection of N2-fixation in association with *S. droebachiensis* (Guerinot and Patriquin 1981a). Although 16S rRNA gene sequences do not *per se* offer any functional information, predicting gene content via inference of genome sequences of phylogenetically related organisms provides a way to gain insights into potential function. Contrary to our hypothesis, predicted genes involved in N2-fixation were not primarily detected during seaweed digestion, but rather in starved sea urchins. This may reflect the higher diversity of the gut microbiome in this treatment. Nevertheless, N2 fixation genes were predicted throughout the experiment in all treatments, indicating that N2-fixing symbionts could be an omnipresent part of the microbiome. To more directly assess the presence of N2-fixing microorganisms, and to identify them taxonomically, we amplified, cloned and sequenced the NifH gene from a selected set of samples of starved, field-captured and kelp-fed sea urchins (*Saccharina* treatment). This demonstrated several lineages of NifH-containing microorganisms within the Gammaproteobacteria, Verrucomicrobia and Bacteroidetes. The majority (10 out of 13) of the clones were related to *Vibrio* sp., which agrees well with the earlier isolation of an N2-fixing strain belonging to this genus from *S. droebachiensis* (Guerinot and Patriquin 1981b).

**Table 2.** Carbon and nitrogen content of the algal food.

| Alga | Mean N<br>% | Mean C<br>% | C:N<br>ratio | Days of<br>storage |
| --- | --- | --- | --- | --- |
| *Fucus serratus* | 2.45 | 35.72 | 14.60 | 4 |
| *Palmaria*<br>*palmata* | 2.46 | 37.37 | 15.21 | 9 |
| *Saccharina*<br>*latissima* | 2.62 | 28.80 | 11.00 | 9 |

### Future perspectives

Our feeding experiment represent the first detailed microbiome data for adult *S. droebachiensis* exposed to different diets and therefore provide a baseline for future studies aimed at disentangling the mechanisms of sea urchin seaweed degradation.

Going from intensively grazing on kelp forests to residing on barrens where food is limited, is made possible by the plasticity of the urchin metabolism. When food is available, sea urchins can process a great deal, but when food is scarce, their metabolism decreases making them able to survive on stored energy from their gonads (Russel 1998). We have shown that a diverse intestinal microbiome is another feature of a starved state, possibly offering different metabolic opportunities in anticipation of food. It is plausible that the intestinal microbiome of sea urchins plays an important role in their plasticity, and key symbionts such as *Psychromonas marina*, nitrogen-fixing taxa, as well as symbionts with yet unknown functions may mediate the response of the sea urchin to food availability. Bacterial symbionts likely offer the metabolic tools needed for efficient kelp digestion and thereby determine the resource use efficiency for sea urchins, influencing their growth and reproduction rates. The feeding fronts with high densities of sea urchins observed at the early stages of destructive kelp forest grazing may be a perfect opportunity for horizontal transfer of intestinal symbionts between sea urchins. Transient symbionts that offer favorable metabolic functions for the local conditions could thereby become abundant and determine the trajectory of a grazing event.

Kelp, as well as other seaweed and seagrasses are key producers of blue carbon (Krause-Jensen *et al*. 2018) which makes their grazers, such as sea urchins, major players in coastal carbon sequestration (Wernberg and Filbee-Dexter 2018). By extension, a dominant symbiont such as *Psychromonas marina*, which likely plays an important role in urchin food degradation, can also have a disproportional influence on carbon cycling. A combination of field experiments to determine the identity and functional potential of symbionts *in situ* during active grazing events, and controlled laboratory experiments involving inoculation of sea urchins with specific symbionts and assessing efficiency of kelp digestion could offer a way forward to gain understanding of host-microbe interactions and organic carbon degradation in the sea urchin microbiome.

In addition to the ecological impact of sea urchin grazing on the marine environment, sea urchins are economically important as a seafood delicacy (Reynolds and Wilen 2000) and make intriguing model systems to study invertebrate evolution (Biermann, Kessing and Palumbi 2003). Their interactions with bacteria, whether they are mutualistic symbionts, or antagonistic pathogens are central to any effort to understand sea urchin biology (Balakirev, Pavlyuchkov and Ayala 2008; Brothers *et al*. 2018; Carrier *et al*. 2021). This study may inform such future investigations, since it highlights important factors which control bacterial diversity and reveals the identity of the dominant symbiont of the green sea urchin.

## Author contributions

MMB conceived the study together with KMN, analyzed the data and wrote the manuscript with contributions from MH and all other coauthors. MH carried out the feeding experiments and contributed to data analysis and interpretation. HW performed the functional prediction and the phylogenetic analyses. SF supervised the experimental work and provided important conceptual input. KMN designed, organized and supervised experimental work and contributed to data interpretation. All authors have read and approved the final version of the manuscript before submission.

## Acknowledgements

We acknowledge Tim Urich for essential infrastructure support and valuable discussions. Sissel Irene Brubak assisted in DNA extraction and Fabian Schäfer assisted in cloning and sequencing of the NifH gene.

## Data Availability Statement

Raw amplicon sequence data has been uploaded to the European Nucleotide Archive under the accession number PRJEB54963. NifH sequences have been submitted to GenBank under accession numbers OP380433-OP380446. Processed data, including ASV tables, taxonomic classification of ASVs and associated metadata will be made available on ResearchGate upon publication (*DOI will be inserted here*).

